# Pulse labeling reveals the tail end of protein folding by proteome profiling

**DOI:** 10.1101/2021.03.28.437442

**Authors:** Mang Zhu, Erich R. Kuechler, Nikolay Stoynov, Joerg Gsponer, Thibault Mayor

## Abstract

Accurate and efficient folding of nascent protein sequences into their native state requires support from the protein homeostasis network. Herein we probed which newly translated proteins are less thermostable to infer which polypeptides require more time to fold within the proteome. Specifically, we determined which of these proteins were more susceptible to misfolding and aggregation under heat stress using pulse SILAC coupled mass spectrometry. These proteins are abundant, short, and highly structured. Notably these proteins display a tendency to form β-sheet structures, a configuration which typically requires more time for folding, and were enriched for Hsp70/Ssb and TRiC/CCT binding motifs, suggesting a higher demand for chaperone-assisted folding. These polypeptides were also more often components of stable protein complexes in comparison to other proteins. All evidence combined suggests that a specific subset of newly translated proteins requires more time following synthesis to reach a thermostable native state in the cell.

## Introduction

Cells have developed an intricate protein homeostasis network to maintain a functional proteome. Homeostasis is constantly challenged by external stresses and internally by unwanted mutations, error during transcription, mRNA processing and translation, as well as a decreased capability through aging (Morimoto and Cuervo, 2014). The result of such challenges frequently culminate in the misfolding of proteins, which can often lead to protein aggregation and potentially deleterious effect in cells (Cooper et al., 2006; Olzscha et al., 2011; Park et al., 2013). Paramount in protein homeostasis is to ensure that each polypeptide reaches, and maintains, its thermostable native state following ribosomal translation.

Folding newly synthesized proteins in a crowded cellular space presents a challenge. Unfolded polypeptides have to potentially explore a large ensemble of conformations, while adopting multiple folding intermediates, to reach their native fold; all the while nonspecific interactions of their exposed linear sequence would interfere with this process (Bryngelson et al., 1995; Ellis and Minton, 2006; Gershenson et al., 2014; Hartl et al., 2011). To cope with such challenge, eukaryotic cells have developed multiple strategies. Translation rate can be slowed for proper co-translational folding (Li et al., 2012; Stein et al., 2019; Zhang et al., 2009) and the absence of co-translational folding leads to protein misfolding (Chang et al., 2005). Co-translational protein folding can be initiated with the formation of alpha-helices within the exit tunnel of the ribosome (Lu and Deutsch, 2005; Ziv et al., 2005) and is frequently aided by ribosome binding chaperones such as Nascent Polypeptide-Associated Complex (NAC) and ribosome-associated complex (RAC). NAC directly binds to the newly synthesized polypeptides to prevent erroneous interactions with other proteins, premature degradation, or mis-localization while RAC recruits Hsp70/Ssb to assist protein folding by preventing the aggregation of newly synthesized proteins (Deuerling et al., 2019; Duttler et al., 2013; Wiedmann et al., 1994). Notably, exposed stretches of hydrophobic regions during translation are protected by Hsp70 chaperones to prevent nonspecific interactions. Understanding of the chaperone network and how NAC, RAC, cytosolic Hsp70/Ssb1 and TRiC/CCT chaperonin assist protein folding co-translationally has been deepened by many recent studies (Döring et al., 2017; Shen et al., 2019; Shiber et al., 2018; Stein et al., 2019; Willmund et al., 2013). However, there is a need to better decipher how newly synthesized proteins are further handled by protein homeostasis network following the release of polypeptides from the ribosome.

Protein folding is typically an extremely fast process, occurring within a few micro-seconds for small domains or proteins (Kaiser and Liu, 2018; Kubelka et al., 2004). However, upon translation of the nascent chain only a portion of the polypeptide can fold at a given time as the synthesis rate is typically slower than the folding rate. As well, the presence of chaperone proteins can slow down the process (Stein et al., 2019) and some polypeptides may only reach their thermostable native state once they are correctly localized in the cell. Whereas assembly of certain protein complexes has recently been shown to occur co-translationally (Shiber et al., 2018; Stein et al., 2019), many large macro-assemblies presumably rely on complex multi-step processes. For instance, ribosomal proteins are trafficked to the nucleolus where they are assembled into preribosomes with the help of assembly factors (Woolford and Baserga, 2013) and unassembled ribosomal subunits are rapidly cleared from the cell to prevent their accumulation (Nguyen et al., 2017; Sung et al., 2016; Yanagitani et al., 2017). Therefore, the extent of which polypeptides can more rapidly reach their thermostable state following translation in comparison to other proteins remains elusive.

Here, we address this gap in knowledge by systemically assessing which newly translated proteins in *Saccharomyces cerevisiae* are not thermostable using a proteomic approach. We find that newly synthesized proteins with increased “pelletability” (i.e. the ability to be enriched in the pellet fraction after centrifugation) following heat shock are shorter and more structured than other proteins in the cell. They often take part in larger protein complexes and are involved in metabolic enzyme activity. Our study provides important new insights into the protein folding landscape in the cell by revealing which newly translated proteins are more susceptible to destabilizing stresses such as heat shock.

## Results

### A subpopulation of proteins is more susceptible to aggregation when newly translated

We sought to determine whether a cluster of newly translated proteins may be more prone to aggregation upon heat-induced misfolding in comparison to other polypeptides. Notably, we hypothesized that a cohort of proteins with distinct features may require more time to fold and reach their native state or be incorporated in complexes. We, therefore, used a pulse SILAC approach (Experiment 1) to identify newly synthesized polypeptides that display a higher propensity to aggregate (i.e., be enriched in the pellet fraction) following heat shock in *Saccharomyces cerevisiae*.

First, we labeled long-lived proteins with light SILAC media in log-phase growing yeast cells incubated at 25°C. We then pulse labelled cells with heavy SILAC media for 15 minutes followed by a 20 minutes heat-shock at 45°C (Fig 1A). Total cell lysate (TCL) and the pellet fractions were collected after centrifugation and then analyzed by bottom-up mass spectrometry. We quantified 3,410 proteins in four biological replicates, including 2,610 proteins quantified in at least three experiments in both fractions (Fig S1A; Table S1). Light to heavy ratios showed strong correlations in pairwise comparisons for samples derived from the same fraction (Fig S1B). Light to heavy ratios in the TCL were normally distributed with a median 10.89% light SILAC labeling (Fig S1C), which is expected for a 15-minutes labeling and 150 minutes cell doubling time. To determine what fraction of newly translated proteins entered the pellet after heat shock, we normalized the light to heavy ratios using the TCL ratios for the 2,610 proteins quantified in at least three out of the four replicates (Fig 1B). In this analysis, proteins isolated in the pellet fraction that consist of long-lived species should have TCL-normalized values near zero in the log scale (i.e., they display similar light to heavy ratios in both fractions). In contrast, proteins that are more represented as newly synthesized polypeptides because they are less thermostable in the pellet fraction should have positive values. We found that 574 proteins were significantly (p-value < 0.05) enriched as newly translated proteins over a two-fold enrichment in the pellet fraction (Fig 1C; Table S1). For instance, the Rpl12 ribosomal protein and the Tdh3 glyceraldehyde-3-phosphate dehydrogenase protein displayed an over 5-fold enrichment (Fig S1D). In addition, 40 proteins appeared to be significantly depleted as newly synthesized proteins in the pellet fraction (Table S1). We found that these depleted proteins have a much higher proportion of heavy labeled peptides in the TCL (i.e., newly synthesized) due to potentially higher translation or turnover rates (Fig S1C). These results show that there is a large cohort of proteins enriched as newly synthesized in the pellet fraction upon heat shock, whereas the majority of newly synthesized polypeptides are not specifically affected by the stress.

**Fig 1.**
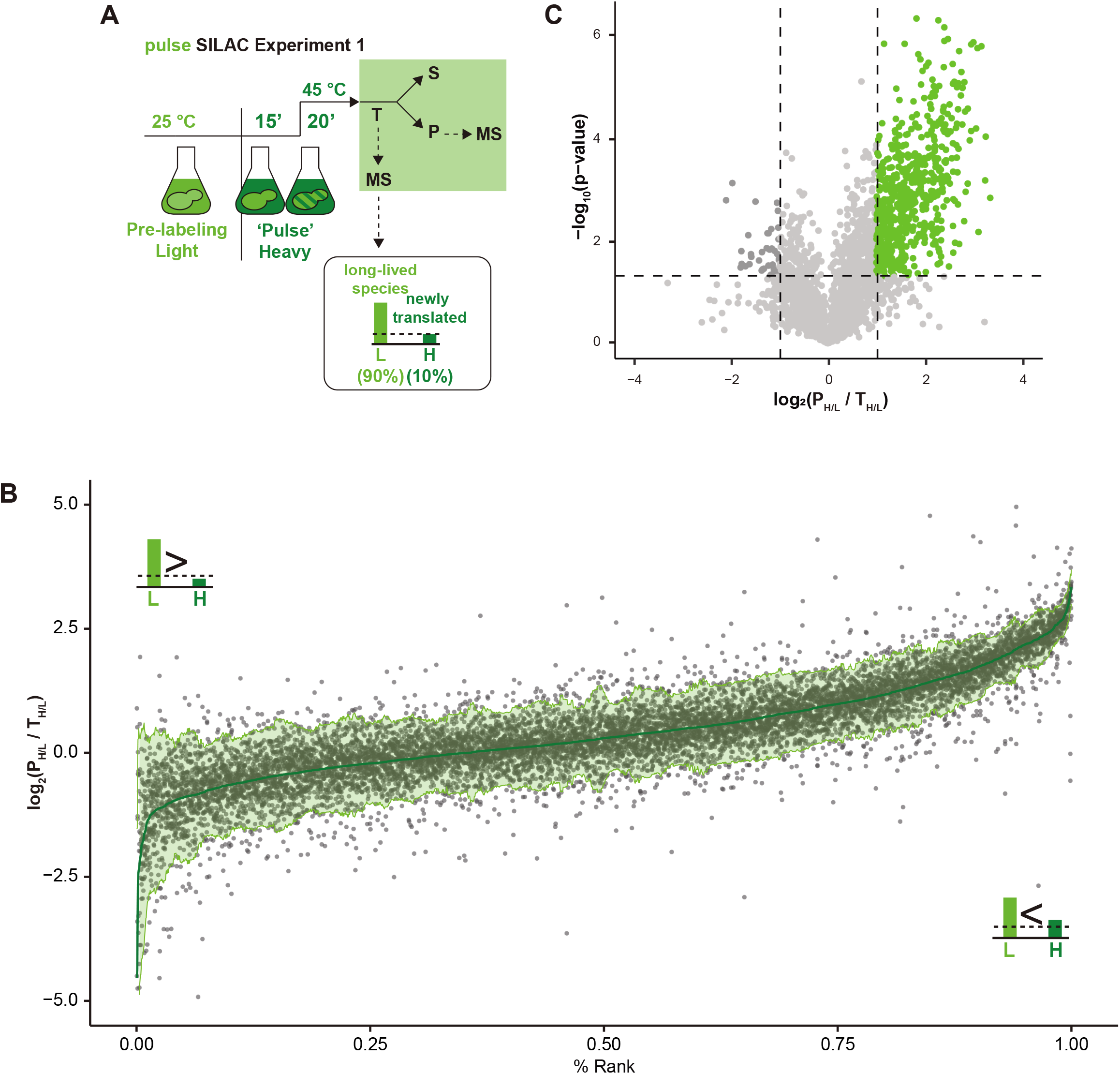
Identification of newly translated proteins that aggregate upon heat stress by pulse SILAC. (A) Schematic of Experiment 1 in which proteins are pre-labelled in light (L) SILAC media prior to a pulse labeling in heavy (H) media and heat shock (S: supernatant fraction, P: pellet fraction, T: total cell lysate). (B) The graph displays the log_2_ values of the normalized SILAC ratios that compare the H/L ratios of the proteins in the pellet fraction (P) over the H/L ratios in the total cell lysate (T). Proteins are ranked based on the averaged log_2_ ratios (green), each gray dot represents one of four replicates if quantified, and the light green shade marks the two standard deviation below and above the mean which is smoothed over a sliding window length of 50. (C) Volcano plot of log_2_ values of the normalized SILAC ratios plotted against the -log_10_ of *t*-test p-values. The dotted lines indicate proteins that have a *t*-test *p*-value < 0.05 and at least a 2-fold change.

### The aggregation of newly translated proteins is not caused by heat-induced translation stalling

We wanted to determine whether the over-represented newly synthesized polypeptides in the pellet might correspond to partially translated proteins that would misfold and aggregate, contributing to an increase of pelletability after heat shock (Fig 2A). However, we found no evidence for a bias toward N-terminal peptides or for higher proportion of light SILAC labeling in the N-terminal regions among the 574 proteins (Fig 2B). Similarly, we did not observe any SILAC labeling bias in the proteins we quantified in the TCL (Fig S2A). As such, we conclude that the possible heat shock induced premature termination of translation does not contribute to the accumulation of newly synthesized polypeptides in the pellet fraction and does not seemingly influence the observed pulse SILAC ratios at the proteome level.

**Fig 2.**
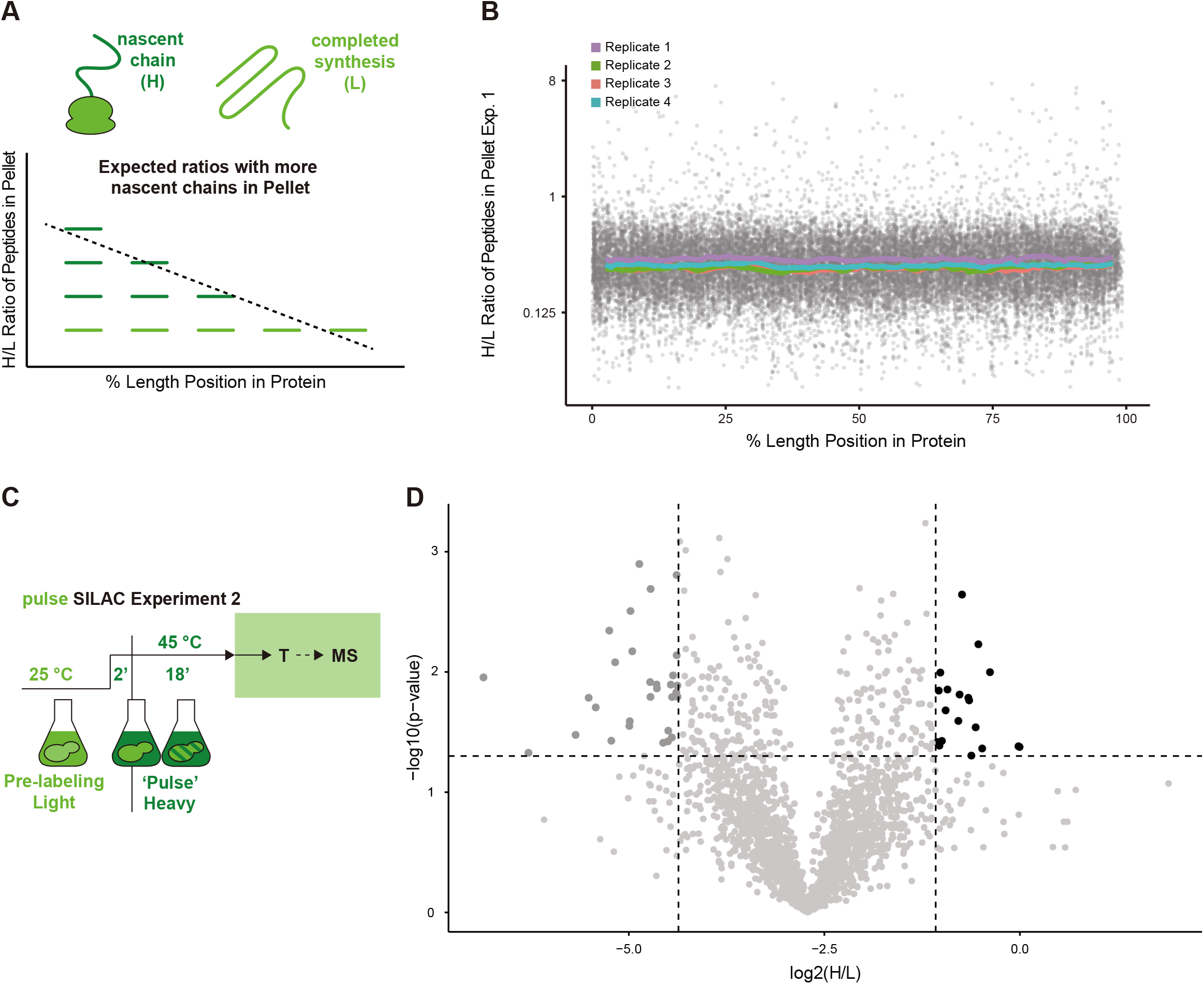
Identification of protein translated during heat shock. (A) Illustration of how partially translated proteins would lead to a N-terminal peptides bias. (B) The graph shows the H/L ratios of peptides of proteins quantified in the pellet fraction in Experiment 1 (y axis) that are positioned based on their location in the protein (x axis). Gray points show individual measurement from all replicates. Colored lines show rolling median ratios with a 5% window and 1% increment of each replicate. (C) Schematic of Experiment 2 in which proteins are pre-labelled in light (L) SILAC media prior to a two minutes heat shock followed by pulse labeling in heavy (H) media (T: total cell lysate). (D) Volcano plot of log_2_ H/L SILAC ratios values plotted against the -log_10_ of *t*-test p-values. The dotted lines indicate proteins that have a *t*-test *p*-value < 0.05 and a labeling over two mean absolute deviation (MAD).

Next, we sought to verify that the pulse SILAC results are not influenced by changes in ribosomal translation that could occur during heat shock. Translation is thought to be inhibited upon acute heat stress (McKenzie et al., 1975; Shalgi et al., 2013). However, we cannot exclude that cap-independent translation mechanisms may still allow some proteins to be synthesized, such as those involving Gis2 (Sammons et al., 2011). Therefore, we carried out a second pulse SILAC experiment where the light SILAC labelled culture was pulsed label with heavy SILAC two minutes after the initiation of the heat shock and continued heat shock for another 18 minutes (Fig 2C; Experiment 2). Cells were collected and lysed for the mass spectrometry analysis of the total cell lysates of three biological replicates. We quantified 2,138 proteins in all three biological replicates that show mostly low level of heavy SILAC labeling (Fig S2B,C; Table S2). We only identified 19 proteins that were significantly more labelled (p values < 0.05) over two mean absolute deviations (MADs) (Fig 2D). Importantly, none of the 19 proteins were among the 574 proteins identified in Experiment 1 and enriched as newly synthesized proteins in the pellet fraction (Fig S2C). In fact, the proteins enriched for newly synthesized in Experiment 1 displayed a lower heavy labelling than average in Experiment 2 (Fig S2C), indicating that their accumulation in the pellet fraction is not due to increased translation during the stress. Collectively, these results suggest that newly synthesized proteins that accumulate in the pellet in the heat shock conditions mostly correspond to polypeptides fully translated prior to the stress.

### Newly synthesized thermolabile proteins are stable when matured

We recently identified a cohort of 86 heat stress granule proteins that are depleted from the supernatant fraction following heat shock (Zhu et al., 2020). When we compared the pulse labeling ratios (Experiment 1) to data from our pervious study, we found that, strikingly, heat stress granule proteins form a distinct cohort, which is constituted of long-lived species that accumulate in the pellet fraction (Fig 3A). From this analysis we can also infer that long-lived proteins of the cohort identified in Experiment 1 mainly remain in the supernatant fraction following heat shock. To validate our conclusions, we fused two proteins, Rpl12b and Ola1, to a tandem fluorescent protein timer with mCherry and super folding (sf) GFP coupled to an inducible copper promoter to track newly translated proteins (Fig 3B). Due to differences in maturation rates of the fluorophores, newly translated proteins with a mCherry-sfGFP tag would display a GFP signal shortly after translation and then would start to display mCherry signal after approximately 30 minutes (Khmelinskii et al., 2012). When tagged with mCherry-sfGFP tag, Rpl12b showed diffused signal for both GFP and mCherry when unstressed (Fig 3C). After incubation at 45°C for 20 minutes, foci formed, indicative of aggregation, with sfGFP but not mCherry. In contrast, the sfGFP signal remained diffused after heat shock in the control experiment, where the tandem timer is expressed alone. These results indicate that while matured Rpl12b species were not recruited to foci, newly translated products were. When we, instead, fused the tandem timer to the Ola1 stress granule protein, the heat induced foci could be seen in both in the sfGFP and mCherry channels (Fig 3C). Taken together, we can conclude that we have identified a unique group of proteins that are not seemingly affected by the heat shock once they “mature” to reach their thermostable conformation but are highly susceptible to aggregation when they are newly synthesized.

**Fig 3.**
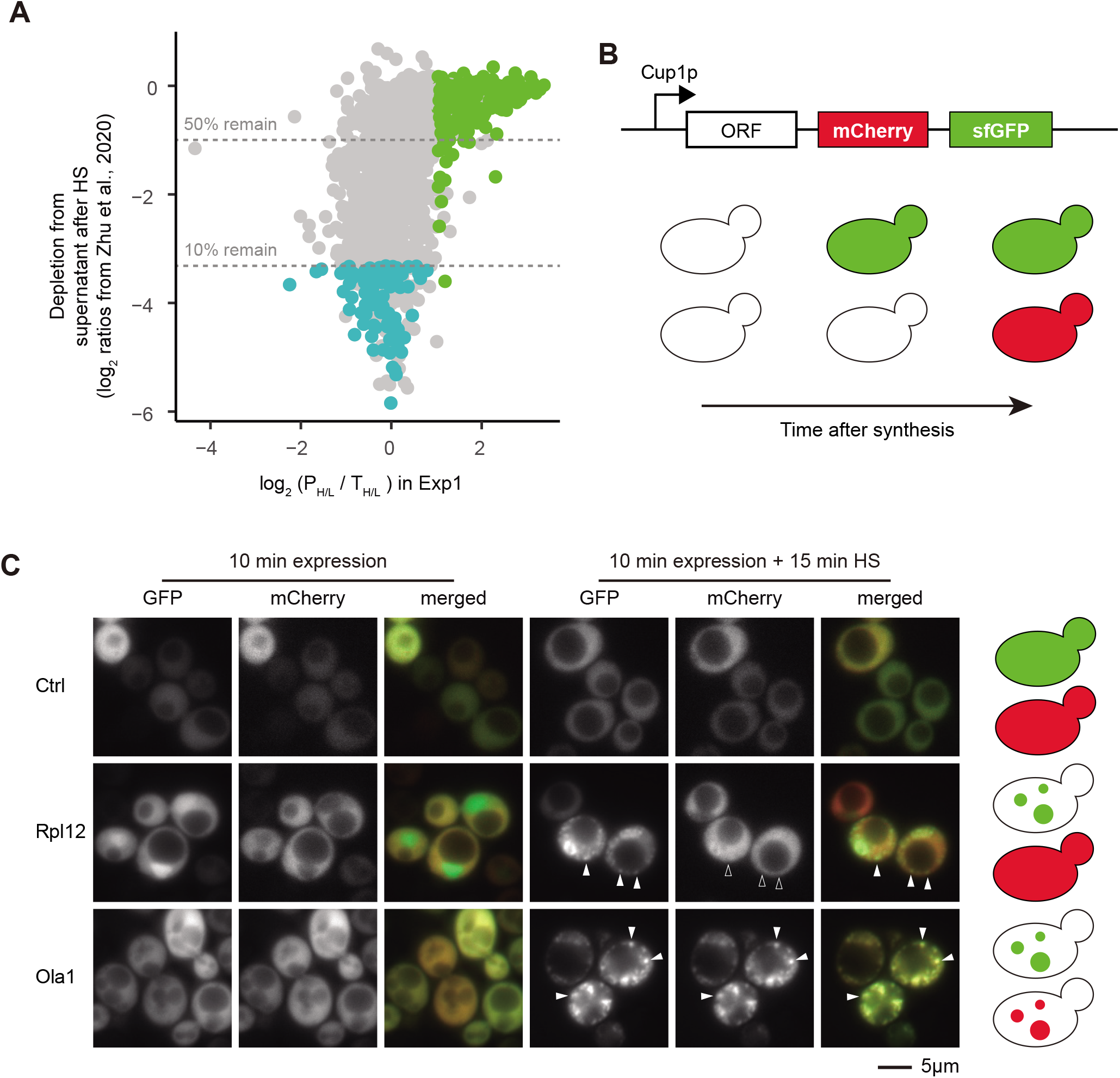
Newly translated proteins enriched in pellet were distinct from stress granule proteins and only form foci in cell as newly translated proteins. (A) Comparison of data from Zhu et al., 2020 and data from Experiment 1. The y-axis displays log_2_ (H/L) of proteins that remain in supernatant after heat shock of heavy-labelled cells (H) vs. control light-labelled cells (L) (proteins with low ratios are depleted from the supernatant fraction) and x-axis displays the log_2_ (P_H/L_/T_H/L_) of proteins in the pellet in Experiment 1. Green dots show newly translated proteins enriched in pellet fraction; Blue dot show stress granule proteins identified in Zhu et al., 2020. (B) Illustration of the tandem fluorescent protein timer construct and expected phenotype of different proteins under heat shock when newly translated. (C) Representative images of cells expressing indicated proteins fused to the tandem fluorescent protein timer. Protein expression was induced for 10 minutes and cells treated with or without 15 minutes heat shock at 45°C. Solid arrowheads show selected foci; hollow arrowheads mark the absence of localization in the selected foci.

### Proteins with distinct characteristics are thermolabile following synthesis

One caveat of Experiment 1 is that it cannot differentiate whether an increase in the pelletability of the newly synthesized proteins is due to heat shock or if these proteins are simply more pelletable when they are newly translated, independent from the stress. To address this issue, we designed a third pulse-SILAC experiment (Experiment 3) to specifically distinguish between newly translated proteins in stress-induced pellet (ntSP) and newly translated proteins constitutively in the pellet (ntCP). We pre-labeled log-phase growing yeast cell in light SILAC media at 25°C and switched to either medium or heavy SILAC media for a pulse labeling of 15 minutes. Medium labeled cells were collected immediately after the pulse labeling while heavy labeled cells were further incubated at 45°C for a 20 minutes heat shock prior to cell lysis, fractionation, and mass spectrometry analysis (Fig 4A). In this particular experiment, ntSP proteins should display a higher heavy to medium SILAC ratios, whereas ntCP should have similar labeling levels. We quantified 2,520 proteins in at least three of the four biological replicates (Table S3), including a large portion of proteins also quantified in Experiment 1 (Fig S4A). When we examined the heavy to medium ratios in the TCL, most proteins displayed similar labeling levels, confirming that translation was shut down due to these heat shock conditions (Fig S4B). Next, we examined proteins that were enriched as newly synthesized protein in the pellet in Experiment 1 (574 proteins) that were also quantified in Experiment 3 (477 proteins, Fig 4B). Indeed, using a two-fold enrichment cut off, we identified 368 ntSP proteins (p-values < 0.05, Table S3). We also identified a cohort of 72 ntCP (p-values > 0.05, less than two-fold enrichment, Table S3). Strikingly, ntSP proteins are significantly enriched for proteins with catalytic activity and involvement in small molecule metabolism (Fig 4C). Many of these proteins also bind to small molecules such as nucleotides and vitamins. When looking at pfam/InterPro domains, proteins with NAD(P)H-binding (29 proteins, p-value = 1.1E-6) and proteasome subunit (11 proteins, p-value = 1.5E-6) were significantly enriched (Table S4). These results suggest that newly synthesized proteins affected by heat shock may share similar structural features that determine whether they are more prone to aggregation.

**Fig 4.**
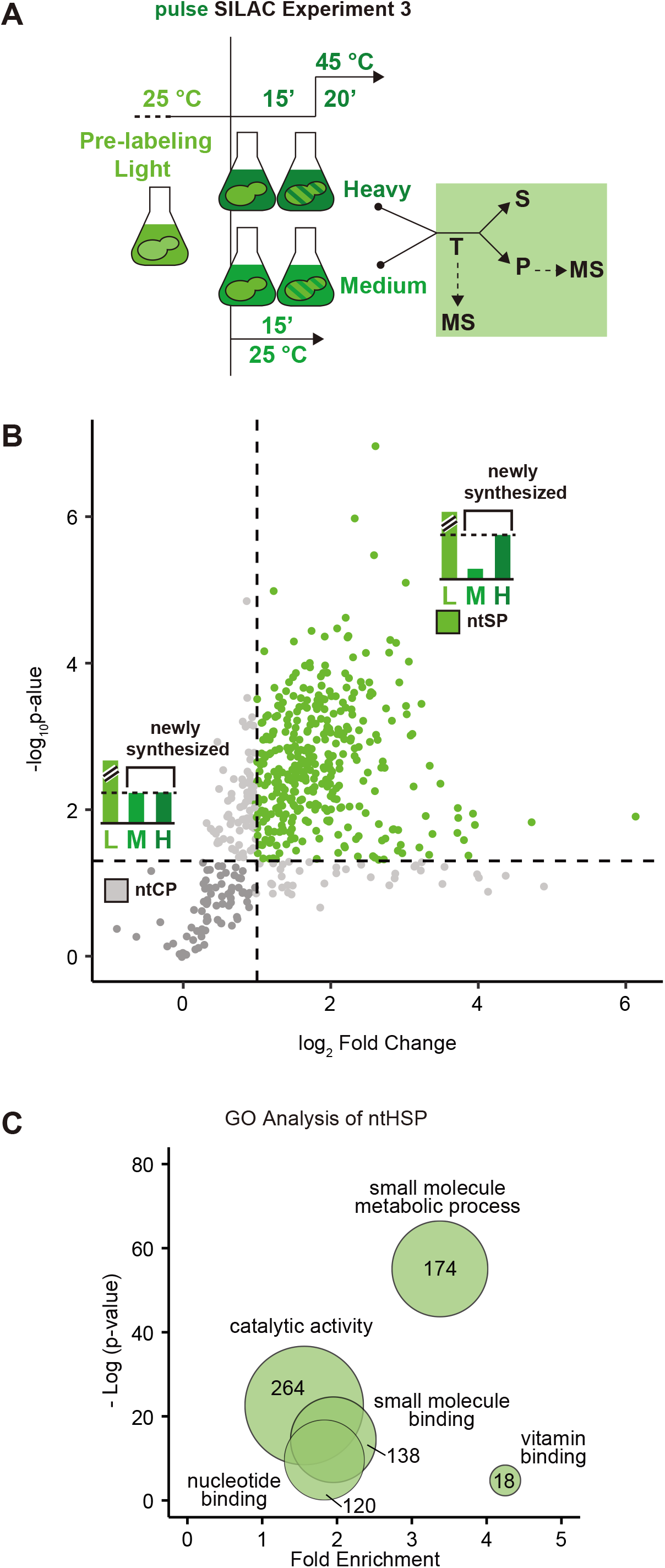
Identification of ntSP and ntCP proteins. (A) Schematic of Experiment 3 in which proteins are pre-labelled in light (L) SILAC media prior to a pulse labeling in heavy (H) media followed by heat shock, or medium (M) media (S: supernatant fraction, P: pellet fraction, T: total cell lysate). (B) Volcano plot of log_2_ values of the normalized SILAC ratios (log_2_ (H_P_/M_P_)/(H_T_/M_T_)) plotted against the -log_10_ of *t*-test p-values. The horizontal dotted line indicates a *t*-test *p*-value of 0.05 and the vertical dotted line indicates a fold change of 2. ntSP proteins are depicted in green and ntCP proteins shown in darker gray. (C) GO analysis of ntSP proteins. Circle size and number indicates the number of proteins within each GO term and placed according to their fold enrichment and significance by Fisher test.

### Newly translated proteins with increase heat-induced pelletability share common features

Mature proteins that have increased pelletability after heat shock were shown to prominently localize in stress granules and these proteins are longer, more disordered and less hydrophobic in comparison to the proteome (Zhu et al., 2020). We recently showed that many of these features can be used to predict the propensity of proteins to be recruited to membraneless compartments such as stress granules (Kuechler et al., 2020). We reason that ntSP proteins represent, instead, proteins that require more time to fold and reach their native state and may therefore display different characteristics.

We performed feature analysis to compare ntSP proteins (n=368) and ntCP proteins (n=72) with all the proteins that did not have over two-fold enrichment in the first experiment (ctrl; n=2,036; Table S1) and the yeast proteome (PME; n = 6,721). The ctrl set was included to ensure that an analyzed feature was not enriched due to our analytical method.

We found that ntSP proteins had significantly grand average of hydropathy (GRAVY) scores and, accordingly, were also predicted to be less solvent exposed and less disordered than both control sets (Fig 5A-C). Additionally, ntSP proteins contain less polar residues than control sets (Fig 5D). One possible explanation is that ntSP proteins have a highly structured core that would explain their absence in the pellet fraction after heat shock once mature. Supporting this observation, a significantly larger fraction of the ntSP proteins’ sequence is, on average, part of pfam domains (Fig 5E). Similarly these proteins display a higher number of pfam domains per residue when compared to controls (Fig S5A) and are enriched for multi-domains (Fig S5B). The length of the polypeptides is unlikely to be driving these results. Whereas these proteins are shorter in sequence length when compared to other proteins quantified in our mass spectrometry analysis, they are longer in comparison to the whole proteome (Fig 5F). Expectedly, ntSP proteins also have significantly less coil regions when compared to the control sets (Fig 5G). Interestingly, we found that ntSP proteins are significantly enriched for β-sheet character (Fig 5H) while the amount of α-helix remains roughly the same (Fig S5C). Polypeptides with β-sheet have been suggested to fold more slowly (Kubelka et al., 2004; Plaxco et al., 1998), and one possibility is that highly structured polypeptides with more β-sheets are less favored for co-translational folding because this secondary structure is often discontinuous in the protein sequence. However, despite a significant enrichment in β-sheet content, ntSP proteins lack a significant elevation in the median number of aggregation-prone stretches to both controls. These regions are calculated using TANGO, a predictor for the propensity of antiparallel β sheet amyloid formation (Fig S5D). These results indicate that there is a subset of proteins that tend to aggregate more following synthesis and heat shock, which share distinct characteristics that is difference from the proteome. Newly synthesized proteins that are in the pellet fraction independently of the stress (ntCP, n= 72), displayed somewhat intermediary features that trended toward ntSP features but that were in most cases not significantly different from controls (Fig 5A-H, S5A, C-E). One caveat is that less proteins are identified as ntCP, which reduces the statistical power of this analysis. Interestingly, ntCP proteins are more disordered than ntSP, as the controls. One possibility is that ntCP share yet another unidentified feature that leads to their aggregation following synthesis in unstressed conditions.

**Fig 5.**
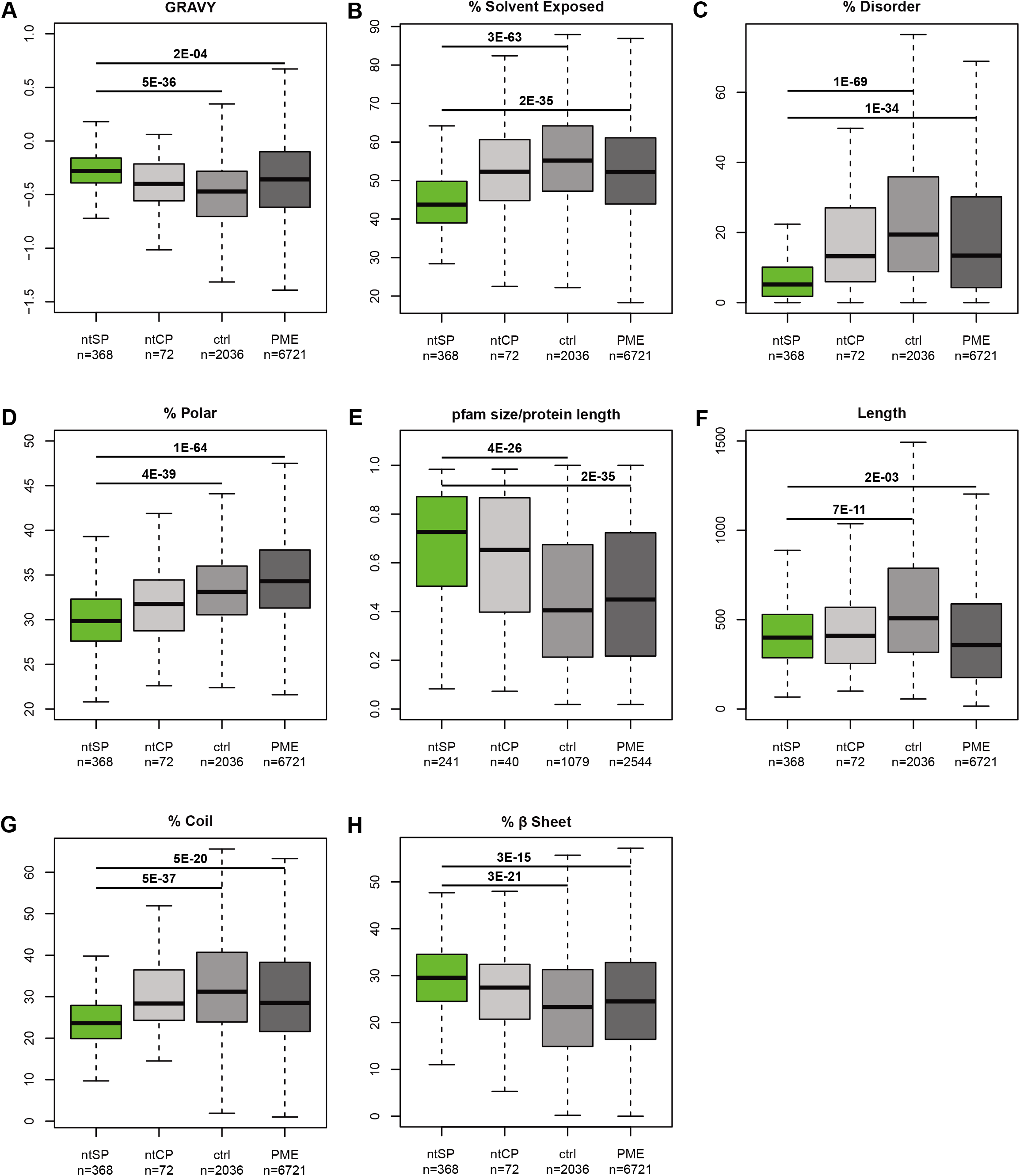
Protein feature analysis of ntSP and ntCP proteins. (A-H) Box plots comparing distributions of ntSP, ntCP, control proteins (ctrl) and proteome (PME). The following analyses are shown: hydrophobicity based on GRAVY score (A), % solvent exposed (B), % disordered (C), % poler (D), the sum of residues within a pfam domain normalized by protein length (E), sequence length in number of amino acids (F), % predicted coil (G), % predicted β sheet (H). p-values (Hochberg adjusted Wilcoxon test) are shown and/or reported in Table S5.

### ntSP are part of larger assemblies with an increased reliance on the protein homeostasis network

We found that ntSP proteins are highly abundant in the cell in comparison to controls (Fig 6A). This result is rather surprising as a higher evolutionary pressure to avoid aggregation of abundant proteins during stress would be expected. For instance, highly expressed proteins were previously shown to evolve at a slower rate, presumably because they have reached optimum translational robustness (Agozzino and Dill, 2018; Drummond et al., 2005). One concern is that the features we observed among the ntSP proteins might be simply driven by their high abundance in the cell. To rule out this possibility, we compared ntSP proteins to the top 20% most abundant proteins from the proteome that show similar abundance than ntSP (Fig S6A). All features previously described remain significantly different between ntSP proteins and high abundant proteins from the proteome (Fig S6B-I). These results indicate that the associated features are unique to ntSP proteins and not simply driven by the high abundance of these proteins in the cell.

**Fig 6.**
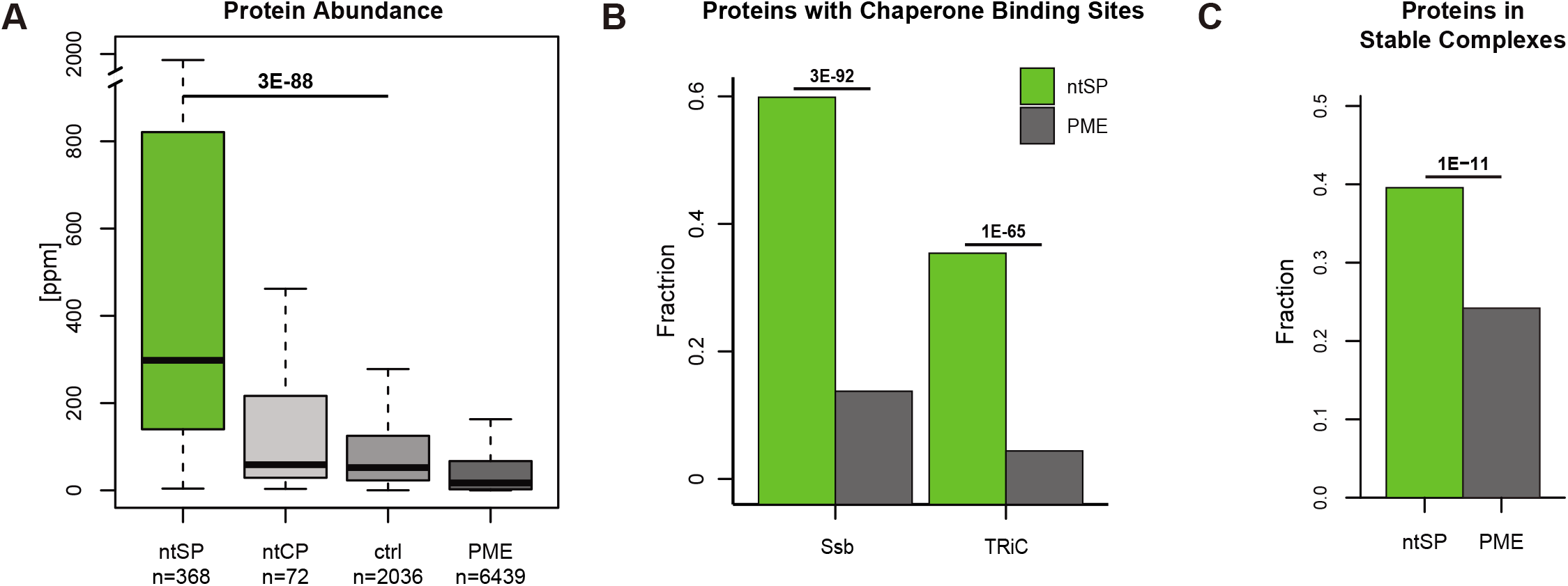
Protein feature analysis of ntSP and ntCP proteins. (A) Box plots comparing protein abundance distributions of ntSP, ntCP, control proteins (ctrl) and proteome (PME). (B) Comparison of proportion of protein with at least one binding site for Ssb (left) and TRiC (right) in ntSP proteins (green) and proteome (PME) based on a study from Dr. Frydman and colleagues (Stein et al., 2019). (C) Comparison of the proportions of proteins in stable complexes within ntSP proteins and proteome (PME). n and p-values (Wilcoxon test in A, Fisher test in B and C) are shown and reported in Table S6.

Next, we sought to determine whether ntSP proteins may be more dependent of specific chaperones for folding. Client proteins of TRiC/CCT were found to be less disordered and have more β-sheet (Stein et al., 2019), reminiscent to some of the features we observed for ntSP. Indeed, and perhaps not surprisingly, we found that ntSP contain a higher proportion of proteins with binding motifs shown to interact with the Ssb Hsp70 chaperone and TRiC/CCT, in comparison to controls (Fig 6B). Therefore, we can conclude that proteins which are reliant on Hsp70 and TRiC/CCT chaperones during translation, generally, require more time to reach their native conformation following their release from the ribosome.

We noticed that many ntSP proteins were also associated to small molecules or nucleic acids (Fig 4C). One possibility is that ntSP are less thermostable in absence of their co-factor. Similarly, we found that ntSP proteins were significantly enriched for polypeptides that are part of a stable protein complex (Fig 6C). Interestingly, we found that proteins that form stable protein complexes in the proteome are generally more disordered, less hydrophobic, and to have more residues that are predicted to be solvent exposed in comparison to other proteins (Fig S6J-L). However, ntSP proteins are depleted for intrinsic protein disorder, perhaps suggesting that this cohort of proteins requires greater assistance from the protein homeostasis network in order to assemble and reach their thermostable state.

## Discussion

Newly translated proteins need to correctly fold to reach their native state. While recent work has deepened our understanding of how nascent polypeptides are handled co-translationally by chaperone proteins, it remained unclear which of these newly translated proteins are handled by the protein homeostasis network following their release from the ribosome remained. Moreover, it is unclear which proteins are more often kinetically trapped during their folding or require more time to reach their thermostable native state. This study provides a systematic survey of the thermostability of newly translated proteins and reveals new insight into newly translated protein homeostasis.

We found a cohort of proteins less thermostable following translation in comparison to other newly synthesized polypeptides, which are referred to as ntSP proteins. We show that the increase of pelletability of these newly translated proteins after heat shock is not due to premature translation termination. We also confirmed that the translation during acute heat shock is largely limited and, thus, has neglectable contribution to the content of newly translated proteins in the pellet fraction.

It is worth noting that ntSP proteins are only enriched in the pellet fraction following their synthesis. When the pelletability of proteins after heat shock is assessed without pulse labelling (Zhu et al., 2020), ntSP proteins are among the most soluble polypeptides in the cell, indicating that the bulk of these proteins is thermostable once mature. In agreement, none of the ntSP proteins are major components of stress granules, which also become highly pelletable following heat shock upon their assembly into a large condensates. Nonetheless, ntSP proteins—but not their folded form—may also be recruited to stress granules and their presence may participate to the formation of the condensates. For instance, chemical inhibition of translation abrogates stress granule formation (Jain et al., 2016; Wallace et al., 2015). More work will be required to assess whether ntSP are in general specifically recruited to stress granules.

Surprisingly, ntSP proteins are highly abundant in the cell. Nonetheless, only a subset of abundant proteins are enriched in the pellet fraction following synthesis, as mature proteins remain in the supernatant. When we compared ntSP proteins to abundant proteins in the proteome, we found that ntSP proteins display distinct features. The presence of abundant proteins in the pellet fraction is perhaps counter-intuitive, as one would expect an evolutionary pressure to reduce their footprint. Nevertheless, the aggregation of these newly synthesized proteins is specific to the heat shock conditions in our experimental framework, which does not typically apply in the natural environment and would therefore not necessary be selected against. In contrast, proteins that are constitutively more prone to aggregation following synthesis (i.e., ntCP) are less abundant, and their aggregation has possibly only a minor impact in cell fitness.

The ntSP proteins show distinct features, such as increased hydrophobicity and having a greater composition of β-sheet structures while predicted to be less disordered than the proteome. Alongside, a larger fraction of ntSP proteins have chaperone binding sites suggesting that these proteins are more dependent on chaperones assistance for folding. Hsp70/Ssb, besides their role in ribosome associated nascent chain folding, were also shown to remain associated with nascent chains after being released the ribosome (Willmund et al., 2013). This association indicates that, perhaps unsurprisingly, protein folding extends far beyond the end of translation. These ntSP proteins may constitute a class of “complex folders”. The thermostability of these folded proteins may come at a cost of longer folding time required after protein synthesis, which can explain their “vulnerability” to heat stress in our pulse-SILAC experiments. In addition, other ntPS protein may aggregate because they remain “orphan” for longer period of time. For instance, ntSP are enriched for proteins that bind with small molecules and are more often part of stable complexes. Indeed, orphaned subunits of stable complexes have been shown to be unstable, as in the case of α-subunit of fatty acid synthetase Fas2 (Scazzari et al., 2015). Therefore, several of these proteins may only reach their thermostable conformation after assembling into larger protein complexes. Similarly, it has also been shown that small-molecule ligands produced by metabolic pathways can stabilize the folded state of proteins on binding by shifting folding equilibria (Fan et al., 1999; Hammarström et al., 2003). Interestingly, ntSP proteins are less disordered in comparison to other proteins whereas proteins in stable complexes are generally more disordered than those not found in complexes. Perhaps, when looking at the isolated sequence, proteins that are apart of complexes must have exposed surfaces for their binding partners to recognize through other features, such as Molecular Recognition Features (MoRFs), that reside within intrinsically disordered regions (IDRs). These IDR would allow unassembled proteins to remain soluble prior to their incorporation. One possibility is that newly translated proteins that are part of larger assemblies and devoid of IDR are more prone to aggregation prior to assembly such as in the case of ntSP proteins.

Our study reveals which newly synthesized proteins remain susceptible to thermal shock for an extended period of time following their release from the ribosome. The discovery that the ntSP set consists of a distinct pool of proteins that share distinct features is instrumental to further understanding of protein homeostasis in the cell.

## Acknowledgments

We thank Dr. Michael Knop for sharing the fluorescent timer protein plasmid, Dr. Eric Jan for support with labeling experiments and members of the Mayor lab for discussions. This work was supported by a grant from the Natural Science and Engineering Research Council of Canada (NSERC; 04248). T.M. is the recipient of Career Awards from the Michael Smith Foundation for Health Research and M.Z. is the recipient of the Alexander Graham Bell Canada Graduate Scholarships-Doctoral and the Killam Scholarship.

## Author Contributions

Experimental design: M.Z., T.M.; Experimental work: M.Z.; Technical support for experimental work: N.S.; Computational analyses: M.Z., E.R.K.; Resources: J.G., T.M.; Writing & editing: M.Z., E.R.K, J.G., T.M.

## Figure legend

**Fig S1.**
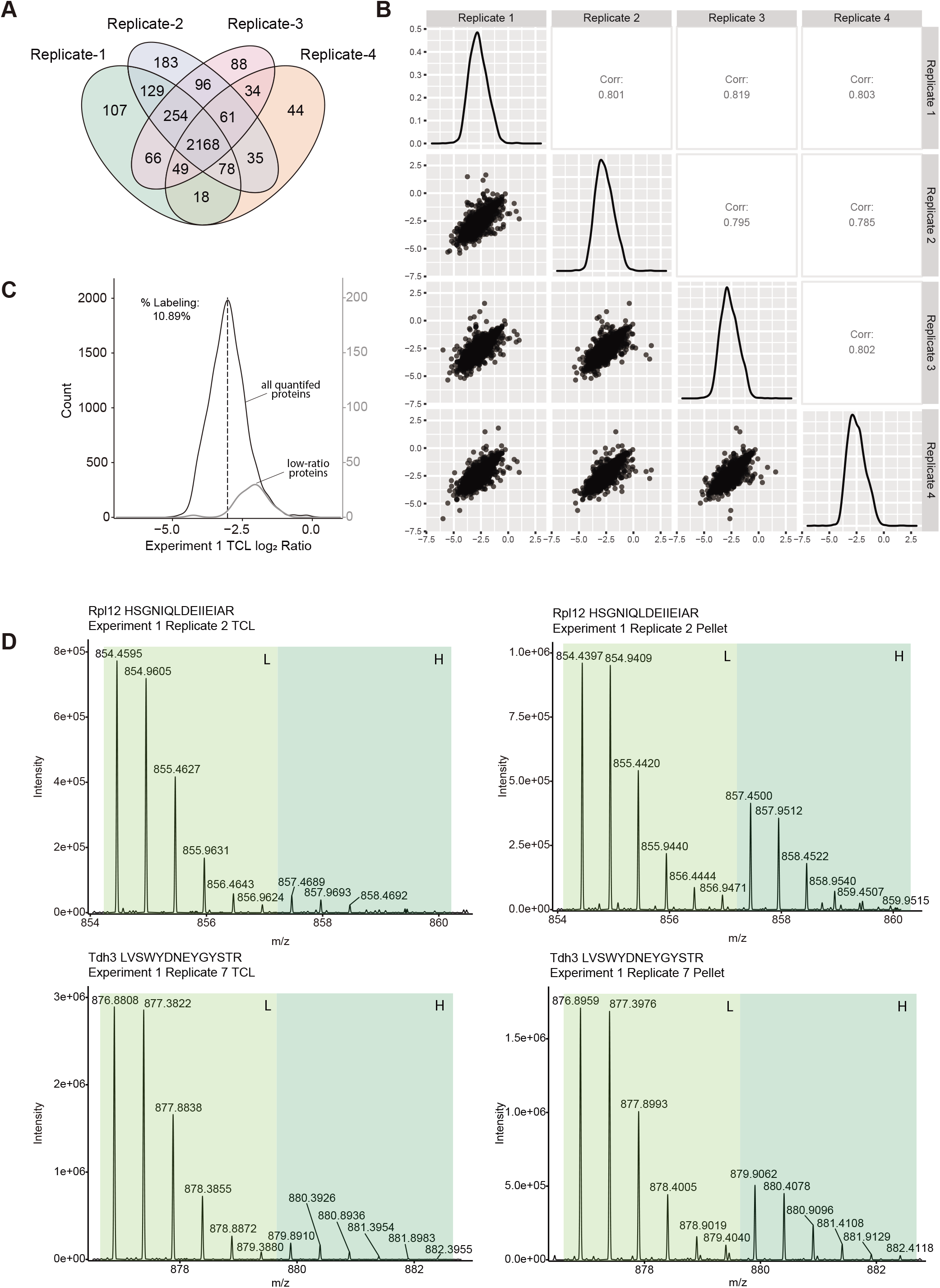
(A) Venn diagram of proteins quantified in each of the four replicates in Experiment 1. (B) Pair-wised comparisons of log_2_ H/L ratios of proteins in the pellet fraction from all four replicates that include scatter plots, distribution plots and Pearson correlation coefficients. (C) Distribution of the averaged log_2_ H/L ratios of proteins in the total cell lysate (TCL) quantified in a least three replicates. The dashed line represents the median value. The grey line shows the TCL ratios of the 40 proteins that were depleted in Figure 1C. (D) Example MS1 spectrums of Rpl12 peptide (HSGNIQLDEIIEIAR) and Tdh3 peptide (LVSWYDNEYGYSTR) from TCL and the pellet fraction.

**Fig S2.**
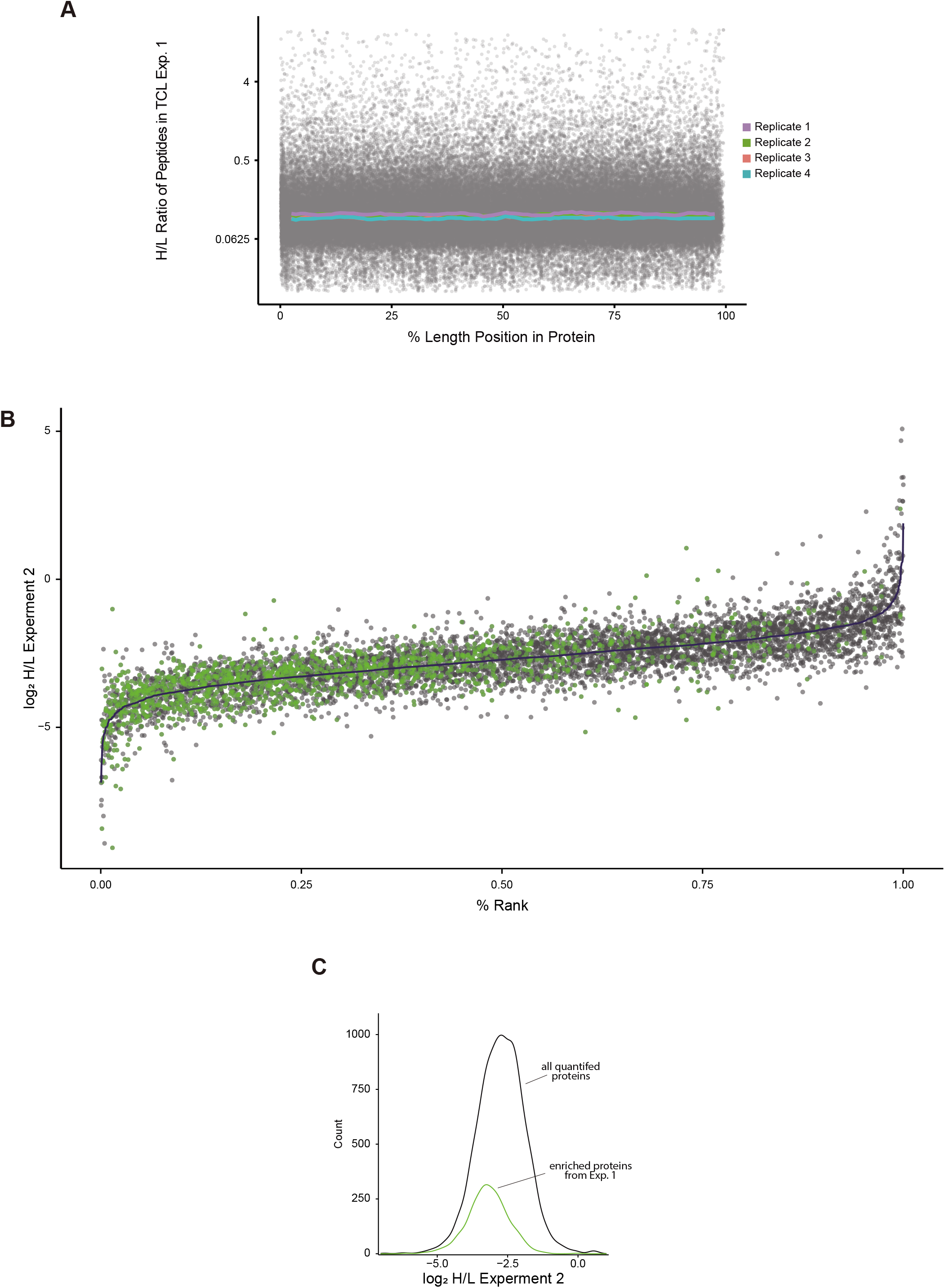
(A) The graph shows the H/L ratios of peptides of proteins quantified in the total cell lysate (TCL) in Experiment 1 (y axis) that are positioned based on their location in the protein (x axis). Gray points show individual measurement from all replicates. Colored lines show rolling median of ratios with a 5% window and 1% increment of each replicate. (B) The graph displays the log_2_ values of the H/L SILAC ratios of the proteins in total cell lysate in Experiment 2. Proteins are ranked based on the averaged log_2_ ratios (dark blue), each gray dot represents one of four replicates if quantified, each green dot represents one of four replicates of proteins enriched as newly synthesized in the pellet fraction from Experiment 1. (C) Distributions of the averaged log_2_ H/L ratios of proteins in the total cell lysate quantified in all three replicates from Experiment 2 (black) and for the 536 proteins enriched as newly synthesized in the pellet fraction in Experiment 1 and quantified in Experiment 2 (green).

**Fig S4.**
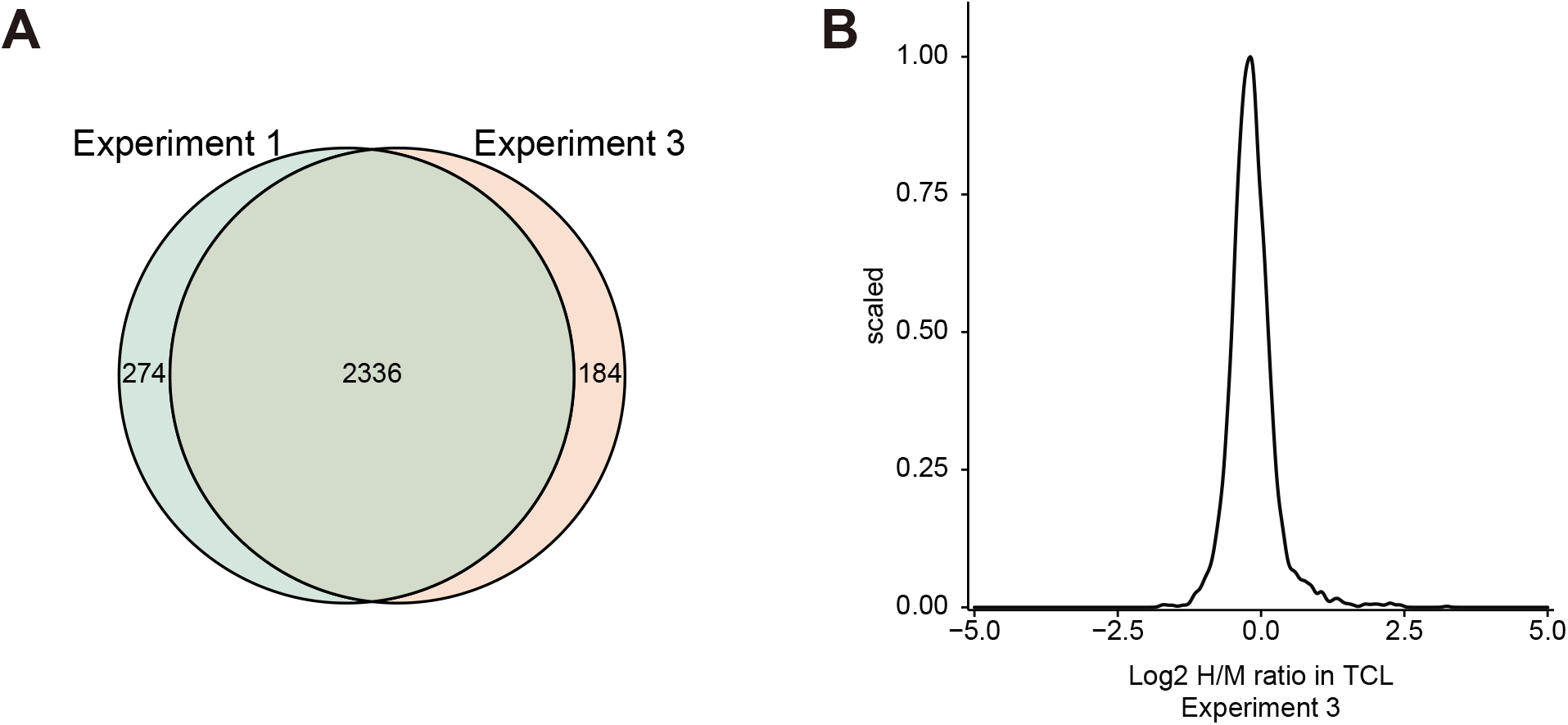
(A) Venn Diagram comparing proteins quantified in Experiment 1 and Experiment 3. (B) TCL log_2_ H/M ratio distribution in Experiment 3.

**Fig S5.**
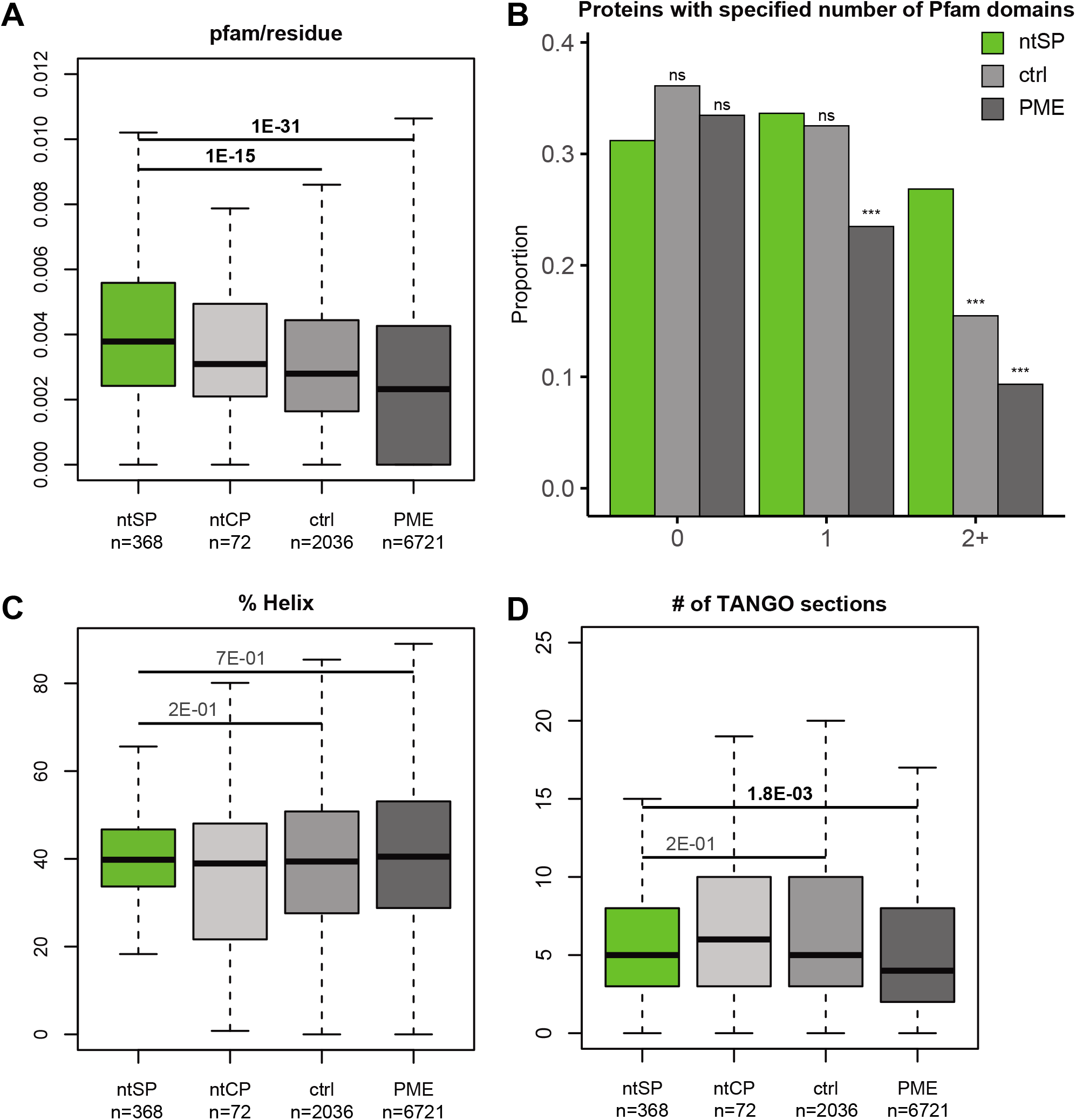
(A, C-D) Box plots comparing distributions of ntSP, ntCP, control proteins (ctrl) and proteome (PME). The following analyses are shown: % charged residues number of pfam domains normalized by protein length (A), % predicted α helix (C), Number of patches of five or more residues predicted to be aggregation-prone by TANGO (D). p-values (Hochberg adjusted Wilcoxon test) are shown and/or reported in Table S5. (B) Proportion of proteins in each group with the indicated numbers of Pfam-assigned domains in ntSP, ctrl and PME. Fisher’s test was applied to compare ntSP with the other two groups: ∗∗∗ indicates p-values <0.005.

**Fig S6.**
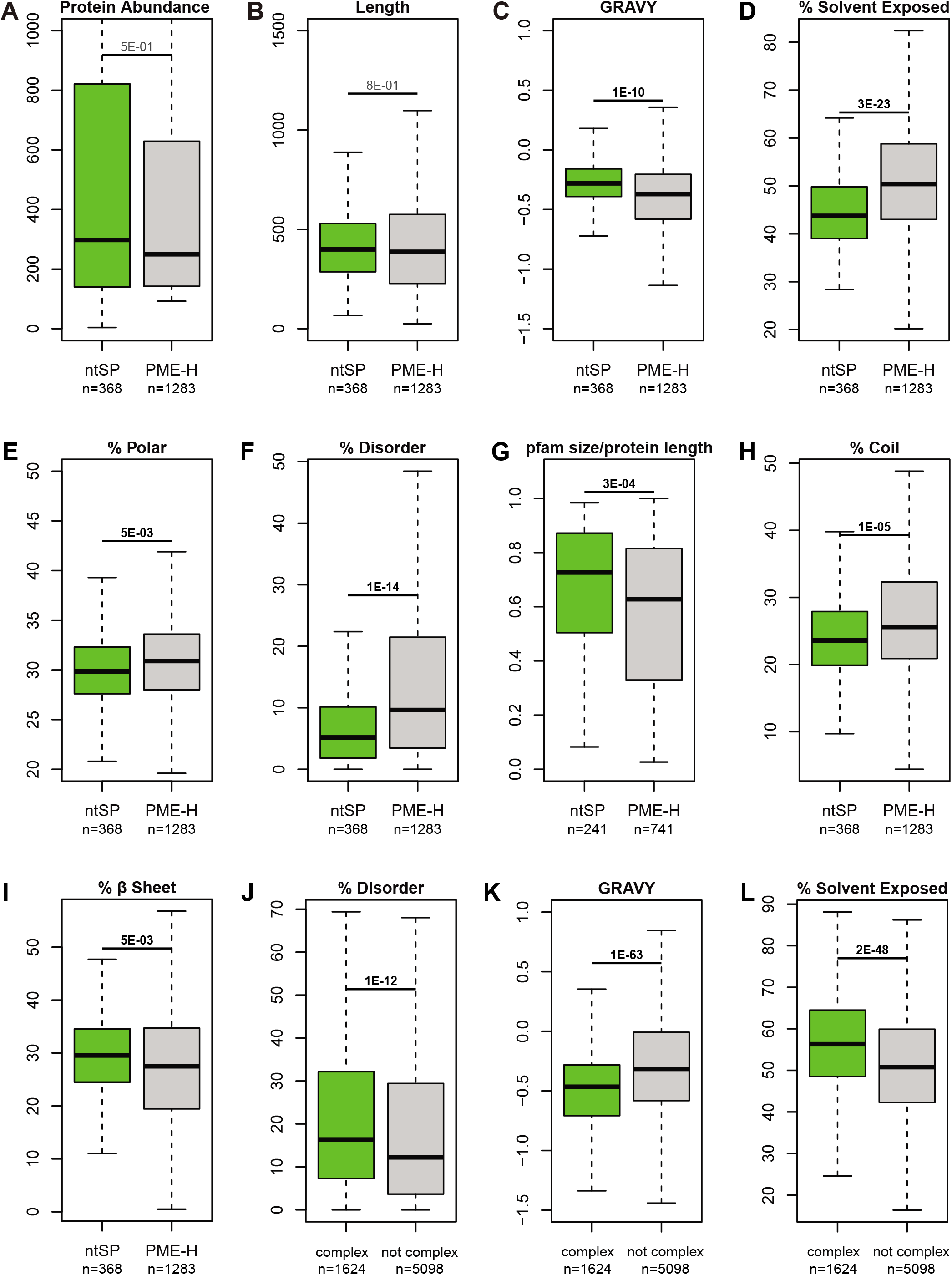
(A-H) Box plots comparing distributions of ntSP and highly abundant proteins in the proteome (PME-H). PME-H correspond to upper quartile The following analyses are shown: protein length in number of amino acids (A), hydrophobicity based on GRAVY score (B), % solvent exposed (C), % poler (D), % disordered (E), pfam domain size normalized by protein length (F), % predicted coil (G), % β predicted sheet (H). n and p-values are shown. (I-K) Box plots comparing distributions of proteins that are known to be part of a stable complex (green) and other proteins (grey). The following analyses are shown: % disordered (I), hydrophobicity based on GRAVY score (J), % solvent exposed (K). n and p-values (Wilcoxon test) are shown and reported in Table S6.

## Tables

**Supplementary Table 1:** Proteins quantified in Experiment 1

**Supplementary Table 2:** Proteins quantified in Experiment 2

**Supplementary Table 3:** Proteins quantified in Experiment 3

**Supplementary Table 4:** Domain Enrichment Analysis of ntSP Proteins

**Supplementary Table 5:** p-values calculated in Figures 5 and S5

**Supplementary Table 6:** p-values calculated in Figures 6 and S6

## Methods

### Yeast strains and plasmids

For SILAC, a modified BY4741 *Saccharomyces cerevisiae* yeast strain was used (YTM1173: *MATa, his3Δ1, leu2Δ0, ura3Δ0, MET15, arg4Δ::KanMX6, lys2Δ0*). pRS316-CUP1p-mCherry-sfGFP-ADH1t (BPM1455) was generated by Gibson assembly of the PCR fragments of pRS316-CUP1p (from BPM1066) and mCherry-sfGFP-ADH1t from pFA6a-mCherry-sfGFP-natNT2 (BPM44, gift from Matthias Meurer). pRS316-CUP1p-RPL12-mCherry-sfGFP-ADH1t (BPM1761) and pRS316-CUP1p-OLA1-mCherry-sfGFP-ADH1t(BPM1762) were generated by PCR of the respective genes from yeast genomic DNA and Gibson assembly into pRS316-CUP1p-mCherry-sfGFP-ADH1t. All tandem fluorescent protein timer plasmids were transformed intoBY4741 cells (*MATa, his3Δ1, leu2Δ0, met15Δ0, ura3Δ0*).

### Yeast culture and Sample preparation

Yeast cells were incubated at 25°C, unless otherwise stated. For the pulse SILAC Experiment 1, cells were grown in light labeled (0.03mg/ml Lys-0, 0.02mg/ml Arg-0) amino acids in synthetic defined (SD) medium with 2% dextrose for at least 7 generations. The equivalent of 80 OD_600_ of cells were taken and washed with heavy labeled SILAC medium (0.03mg/ml D_4_ Lys-4 and 0.02mg/ml ^13^C_6_ Arg-6) twice at room temperature and then resuspend in 100ml heavy SILAC medium and incubated at 25°C for 15 minutes before heat shock for 20 minutes at 45°C in a shaking water bath before cell collection. For pulse SILAC Experiment 2, cells were grown in light labeled (0.03mg/ml Lys-0, 0.02mg/ml Arg-0) amino acids in SD medium with 2% dextrose for at least 7 generations. The equivalent of 80 OD_600_ of cells were taken and washed SD-Lys medium (with no Lys or Arg) twice at room temperature and then resuspend in 100ml SD-Lys medium. The culture was heat shock for 2 minutes at 45°C before the addition of heavy labeled Lys and Arg to a 2x final concentration (0.06mg/ml Lys-4 and 0.04mg/ml Arg-6). The culture was heat shocked for another 18 minutes in a shaking water bath before cell collection. Cells were collected as in Experiment 1. For pulse SILAC Experiment 3, cells were grown in light labeled (0.03mg/ml Lys-0, 0.02mg/ml Arg-0) amino acids in SD medium with 2% dextrose for at least 7 generations. An equivalent of 80 OD_600_ of cells were taken and washed with either medium (0.03mg/ml Lys4 and 0.02mg/ml Arg6) or heavy (0.03mg/ml ^13^C_6_^15^N_2_ Lys-8 and 0.02mg/ml ^13^C_6_^15^N_4_ Arg-10) labeled SILAC media twice at room temperature and then resuspend in 100ml of their respective SILAC medium and incubated at 25°C for 15 minutes. The medium labeled culture was collected after 15 minutes incubation and the heavy labeled culture was heat shock for 20 minutes at 45°C in a shaking water bath before cell collection. All cell harvests were done by centrifugation at 3,220x*g* for 5 minutes at 4°C, washed twice with cold 1×TBS (50mM Tris pH 7.5, 150mM NaCl), mixed, and re-suspended in equal volume of 2×Native lysis buffer (200mM Tris pH 7.5, 150mM NaCl, 2mM PMSF, 2×Protease Inhibitor Cocktails (Sigma-Aldrich)). Re-suspended cells were snap-frozen drop by drop in liquid nitrogen. The resulting cells pellets were lysed by cryo-grinding in a Retsch MM400 (4×30Hz for 90sec) with 1.5ml grinding jar (Retsch #014620230) and one 2mm stainless steel grinding ball (Retsch #224550010). The lysates were thawed on ice and further diluted 3 times by adding ice-cold 1×Native lysis buffer with 1% NP40.

Total cell lysates were pre-cleared twice by centrifugation at 1,000×*g* at 4°C for 15min. Pellet fractions were separated from the resulting supernatant by centrifugation at 16,100×*g* at 4°C for 15min. The insoluble protein pellet was washed twice with ice-cold 1×Native lysis buffer containing 1% NP40 and re-solubilized in 1×Laemmli Sample Buffer. Protein concentrations were determined by using BioRad DC(tm) Protein Assay.

### Mass spectrometry

200μg of protein samples from pellet or total cell lysate fractions were cleaned up and trypsin digested using Single-Pot Solid-Phase-enhanced Sample Preparation (SP3) method as previously described (Hughes et al., 2019). The resulting peptides were desalted with high-capacity C18 (Phenomenex) STAGE Tips before an offline high pH reversed-phase chromatography fractionation as previously described (Udeshi et al., 2013). Fractions were collected at 2 minutes intervals. The resulting 40 fractions were pooled in a non-contiguous manner into 12 fractions per SILAC experiment. Each fraction was then dried in a Vacufuge plus (Eppendorf). Prior to mass spectrometry analysis each sample was resuspended in 0.1% formic acid with 2 % acetonitrile in water.

Mass spectrometry spectra were acquired on an Impact II (Bruker) on-line coupled to either an EASY-nLC 1000 (Thermo Scientific) or a nanoElute (Bruker) liquid chromatography (LC). The LC was equipped with a 2-cm-long, 100-μm-inner diameter trap column packed with 5 μm-diameter Aqua C-18 beads (Phenomenex) and a 40-cm-long, 50-μm-inner diameter fused silica analytical column packed with 1.9 μm-diameter Reprosil-Pur C-18-AQ beads (Dr. Maisch) and it was heated to 50°C using tape heater (SRMU020124, Omega.com and in-house build temperature controller). Buffer A consisted of 0.1% aqueous formic acid and 2 % acetonitrile in water, and buffer B consisted of 0.1% formic acid in 90 % acetonitrile Samples were run with a 120 minutes gradient from 10% Buffer B to 17% Buffer B during the first 45 min, then Buffer B was increased to 35% by 90 min and to 80% at 95 min. The scanning range was from m/z 200 to 2000 Th. The Impact II was set to acquire in a data-dependent auto-MS/MS mode with inactive focus fragmenting the 20 most abundant ions (one at the time at 18 Hz rate) after each full-range scan from m/z 200 Th to m/z 2000 Th (at 5 Hz rate). The isolation window for MS/MS was 2 to 3 Th depending on parent ion mass to charge ratio and the collision energy ranged from 23 to 65 eV depending on ion mass and charge (Beck et al., 2015). Parent ions were then excluded from MS/MS for the next 0.3 min and reconsidered if their intensity increased more than 5 times. Singly charged ions were excluded since in ESI mode peptides usually carry multiple charges. Strict active exclusion was applied. Mass accuracy: error of mass measurement is typically within 5 ppm and is not allowed to exceed 10 ppm. The nano ESI source was operated at 1900 V capillary voltage, 0.25 Bar nanoBuster pressure with methanol in the nanoBooster, 3 L/min drying gas and 150°C drying temperature.

All raw proteomics data have been deposited to the ProteomeXchange Consortium through the PRIDE partner repository with the identifier PXD024336 (Deutsch et al., 2016; Vizcaíno et al., 2014). The results were analyzed against SGD_orf_trans_all_20150113 released from the *Saccharomyces* Genome Database (yeastgenome.org) with common contaminants with the latest version of MaxQaunt software when the data was generated (version 1.6.14 for Experiments 1 & 3, version 1.5.7.4 for Experiment 2). The searches were done using the following parameters: multiplicity matching specific SILAC experiment design for Lys and Arg isotope type, trypsin enzyme specificity, fixed modifications - carbamidomethyl, variable modifications - methionine oxidation and N-acetyl peptides with default software instrument specific search settings, plus match-between-runs and re-quantification and an FDR set below 0.01.

### Bioinformatics

Graphs in figures were generated using the R and assembled using Adobe Illustrator (R Core Team, 2018). Computational analysis tools were carried out using in-house Perl scripts and statistical tests were performed in R. In all cases of non-binary data, p-values were obtained using the Mann-Whitney-Wilcoxon test in conjunction with the Benjamini-Hochberg procedure for multiple testing corrections to determine significance thresholds and reduce the false discovery rate (Bauer, 1972). GO analysis was done using gprofiler2 R package (Raudvere et al., 2019). IDRs were determined with DISOPRED3 (Jones and Cozzetto, 2015). Patches of disordered amino acids were identified using a sliding window of 30 amino acids which allowed for up to 5 amino acids to be considered as ‘not disordered’ before truncating the disordered stretch. Protein abundances were used from the integrated yeast proteome on PaxDb 4.0 (Wang et al., 2015). Secondary structure and solvent exposure calculations were performed using the SSpro8 and ACCpro software in the SCRATCH-1D 1.1 package (Magnan and Baldi, 2014). GRAVY score was calculated with in house scripts using previously derived parameters (Kyte and Doolittle, 1982). Aggregation prone patches were determined by TANGO (Linding et al., 2004) software package. Yeast protein complex data was extracted from CYC2008 2.0 (Pu et al., 2008). Hsp70/Ssb and TRiC/CCT binding site data was extracted from the study done by Stein and colleague (Stein et al., 2019). Pfam domain enrichment was done using DAVID (Huang et al., 2009a, 2009b).

### Microscopy

For epifluorescence microscopy, cells were grown to mid-log phase (∼OD_600_ = 0.6) in SD medium with low fluorescence yeast nitrogen base (LoFlo, Formedium) and 2% dextrose. Expression of the protein of interest was induced by addition of CuSO_4_ to the final concentration of 100µM for 10 minutes. 1ml of cultures were heat shocked in a thermomixer at 45°C for 15 minutes. Cells were subsequently collected by centrifugation in a microfuge at 6,000x*g* for 30 seconds. Live cells were resuspended in growth media before imaging with a Zeiss AxioObserver Z1 equipped with a 470nm and 590nm Colibri light sources and a 63× 1.4 NA oil immersion DIC objective. Images were acquired and processed with the Zen 3.0 blue edition software.

